# MS2Query: Reliable and Scalable MS^2^ Mass Spectral-based Analogue Search

**DOI:** 10.1101/2022.07.22.501125

**Authors:** Niek F. de Jonge, Joris R. Louwen, Elena Chekmeneva, Stephane Camuzeaux, Femke J. Vermeir, Robert S. Jansen, Florian Huber, Justin J.J. van der Hooft

**Author notes:** Shared last authors (Florian Huber, Justin J.J. van der Hooft). **Author contribution statement:** NFJ wrote the main code for MS2Query, did the benchmarking, and wrote the first draft of the paper. FH did many code reviews and wrote part of the code of MS2Query. JRL built the first prototype for MS2Query. JJJvdH, FH and JRL extensively reviewed and edited the manuscript. EC and SC shared the data and manually validated the results for case study 1-3. FJV and RSJ collected the data and manually validated the results for case study 4. FH and JJJvdH supervised the work. All authors read and approved the manuscript.

## Abstract

Metabolomics-driven discoveries of biological samples remain hampered by the grand challenge of metabolite annotation and identification. Only few metabolites have an annotated spectrum in spectral libraries; hence, searching only for exact library matches generally returns a few hits. An attractive alternative is searching for so-called analogues as a starting point for structural annotations; analogues are library molecules which are not exact matches, but display a high chemical similarity. However, current analogue search implementations are not yet very reliable and relatively slow. Here, we present MS2Query, a machine learning-based tool that integrates mass spectral embedding-based chemical similarity predictors (Spec2Vec and MS2Deepscore) as well as detected precursor masses to rank potential analogues and exact matches. Benchmarking MS2Query on reference mass spectra and experimental case studies demonstrates an improved reliability and scalability. Thereby, MS2Query offers exciting opportunities for further increasing the annotation rate of complex metabolite mixtures and for discovering new biology.

## Introduction

Wide-screen untargeted metabolomics applications are increasingly used to understand complex metabolite mixtures. To boost the metabolite structure annotation rate, mass spectrometry fragmentation approaches are a key source of information in the field of metabolomics^1^. Many improvements have been made in automatically elucidating molecular structure from mass spectrometry fragmentation spectra (also referred to as MS/MS or MS^2^ spectra)^2^. However, it remains very challenging to reliably determine structures based on MS^2^ spectra^3^. Currently, three main types of approaches to determine molecular structures from MS^2^ spectra exist: matching against annotated mass spectral library spectra^4-9^, by using fragmentation trees^10-12^, or by using in silico methods to match against structural libraries^13-15^. However, all these approaches still have important limitations.

One inherent limiting factor of mass spectral library matching is that annotated spectra for only a fraction of the chemical space are known. For example, the GNPS^16^ public mass spectral libraries contain about 2.5% of known natural products^17^. When searching for exact matches, this typically results in finding a few exact spectral matches (with corresponding molecular masses) in a given sample^18^. To overcome this limitation several methods try to search larger structure databases like Pubchem^19^ for potential matches. These methods typically rely on first predicting spectra from structures by using in silico fragmentation, followed by comparing MS^2^ spectra to these predicted spectra^13-15^. Even though these methods are promising, they are still far from perfect at predicting in silico fragmentation, especially for larger molecules such as complex secondary metabolites or lipid-like molecules. Other methods try to retrieve information directly from the MS^2^ spectra without relying on library databases by creating fragmentation trees. Fragmentation trees have been used to predict molecular formulas^10^, used for matching against structural databases^10, 12^, for predicting molecular fingerprints^20^ and recently for completely novel predictions of structures from MS^2^ spectra^11^. These methods show excellent results for smaller metabolites of <400 Da, however for larger metabolites these approaches are still not fully reliable in returning correct elemental formulas and candidate structures. Besides that, the computation time to determine the fragmentation trees also increases manifold. Natural mixtures typically contain considerable amounts of larger metabolites (>800 Da), and this thus poses challenges on the mass spectral interpretation.

A different approach to increase the percentage of spectra for which chemical information can be retrieved is by searching for analogues instead of exact matches. This approach also relies on annotated mass spectral libraries, but aims at finding molecules that are chemically similar, without the need for them to be identical. Current tools able to perform analogue searches often rely on a (modified) cosine score to predict chemical similarity^4, 9, 21^. However, a limitation of the cosine score (and its derivatives) is that small chemical modifications can, and multiple chemical modifications will, often result in a large decrease in mass spectral similarity which limits its ability to serve as a proxy for chemical similarity^22-24^. Recently, two machine learning-based methods were developed that outperform cosine-based scores in predicting chemical similarities from MS^2^ mass spectral pairs; the unsupervised Spec2Vec^23^ and the supervised MS2Deepscore^25^. We hypothesised that their chemical similarity predictions offer great potential for performing a reliable analogue search.

Here we present MS2Query, a tool for rapid large-scale MS^2^ library matching that enables searching both for analogues and exact matches in one run. MS2Query can reliably predict good analogues and exact library matches. We demonstrate that MS2Query is able to find reliable analogues for 35% of the mass spectra during benchmarking with an average Tanimoto score of 0.67 (chemical similarity). This is a substantial improvement compared to the modified cosine score-based method (currently most widely used^4, 9^), which on the same test set resulted in an average Tanimoto score of 0.45 with settings that resulted in a recall of 35% (percentage of query spectra for which a match is predicted). For this benchmarking test set, any exact library matches were removed from the reference library to make sure the best possible match that can be found is an analogue. MS2Query performs especially well for molecules larger than 600 Da: for spectra in a test set in this mass range without any exact library matches, MS2Query predicts an analogue with an average Tanimoto score of 0.85 (high chemical similarity) and a recall of 63%. Besides thorough benchmarking on annotated library spectra, MS2Query was also used for multiple case studies. The higher accuracy of MS2Query offers exciting opportunities for further increasing the annotation rate of complex metabolite mixtures and for discovering new biology. MS2Query is available as a well-tested, open source Python library which grants easy access for researchers and developers.

## Results

### MS2Query combines several machine learning approaches for a more reliable analogue search

The workflow for running MS2Query first uses MS2Deepscore^25^ to calculate spectral similarity scores between all library spectra and a query spectrum. In contrast to existing methods, no preselection on precursor m/z is needed. By using pre-computed MS2Deepscore embeddings for library spectra, this full-library comparison can be computed much faster than existing alternatives (see Speed Performance section). Next, the top 2,000 spectra with the highest MS2Deepscore are selected. MS2Query optimises re-ranking of the best analogue or exact match at the top by using a random forest that combines five features. The random forest predicts a score between 0 and 1 between each library and query mass spectrum. By using a minimum threshold for this score, unreliable matches can be filtered out.

As input for the random forest model, MS2Query uses five different features, calculated between the query spectrum and each of the 2,000 preselected library spectra. These features are Spec2Vec similarity^23^, query precursor m/z, precursor m/z difference, a weighted average MS2Deepscore over 10 chemically similar library molecules, and the average Tanimoto score for these 10 chemically similar library molecules. The random forest model was trained to predict Tanimoto scores (molecular fingerprint based chemical similarity) based on these 5 features. More details about the rationale behind these features can be found in supplementary information S1 and the material and methods.

### Speed performance

Running MS2Query on 5,987 test spectra took 1 hour and 14 minutes (80 spectra per minute) on a normal laptop with an 11th generation Intel Core i5-1135G7 and 16 GB of RAM. The test spectra were matched against a library of 302,514 spectra, without doing any preselection on the precursor m/z difference. An analogue search on the same test set using the matchms implementation^26^ of the Modified cosine score and a preselection on a maximum precursor m/z difference of 100 Da took 9 hours and 24 minutes (10,6 spectra per minute). Note that this would take much longer with a larger maximum precursor m/z difference.

### MS2Query has a high accuracy for analogue searching and exact matching in benchmarking

The performance on finding exact matches and finding analogues was tested separately using two different test sets. The test set for searching for exact matches (‘exact match test set’) contains 3,000 spectra that have at least one spectrum in the library from exactly the same molecule. The test set to test the performance for an analogue search (‘analogues test set’) contains spectra that do not have an exact match to a library spectrum. Thus, for this test set, the best possible match has to be an analogue of the query spectrum.

The performance of MS2Query was compared to MS2Deepscore and the (modified) cosine score for finding analogues and exact matches. For all three methods a minimal threshold can be used to vary the percentage of query spectra for which a match is predicted (recall). For all three methods, the accuracy increases with more stringent thresholds, but the recall decreases. To assess performance for an analogue search, recall is compared to accuracy on the ‘analogues test set’ (Figure 2a). As a metric for accuracy the average Tanimoto score between the test molecules and predicted analogues is used. The Tanimoto score^27^ is a metric for chemical similarity between two molecules, based on chemical fingerprints^28^. Across all recall values, MS2Query has a higher accuracy than comparable search methods relying solely on MS2Deepscore or on the modified cosine score. When aiming for a recall of 35%, the MS2Query threshold for this test set is 0.633 and results in an average Tanimoto score of 0.67 (Figure S3 shows a detailed Tanimoto score distribution).

**Figure 1:**
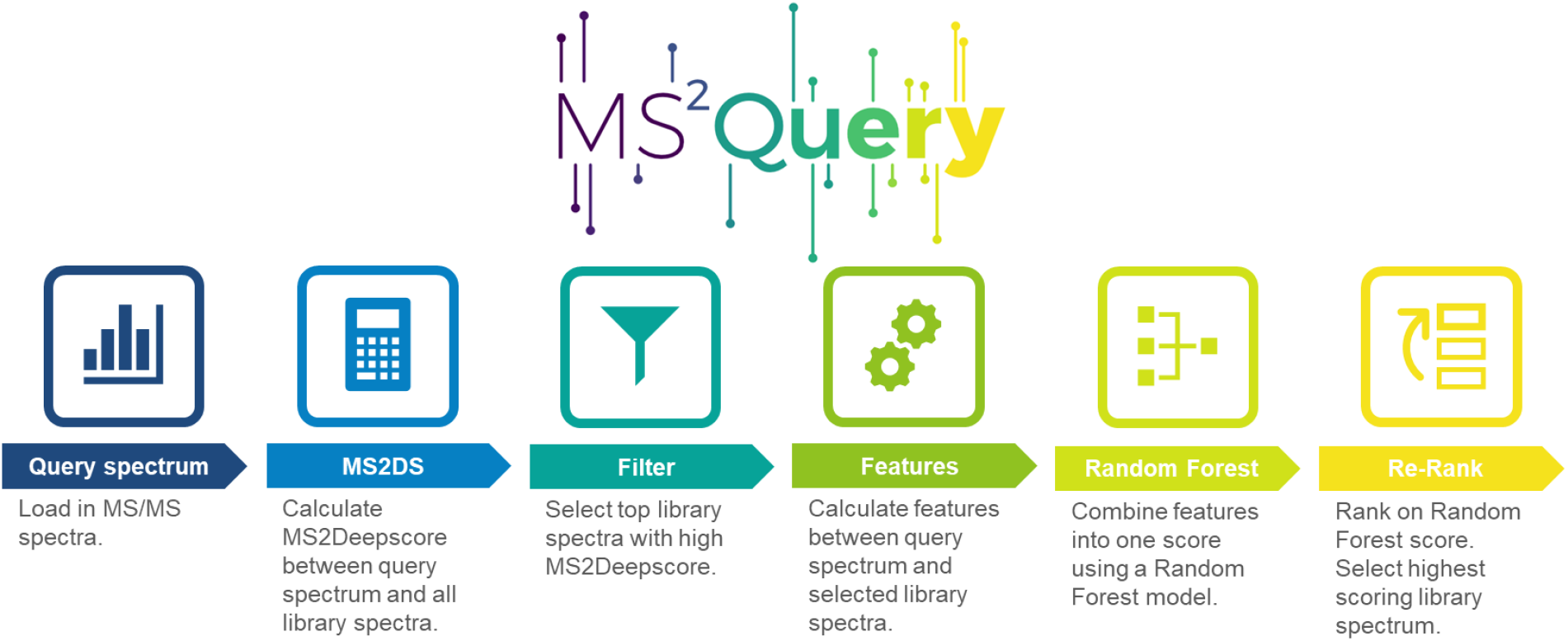
Schematic workflow of MS2Query. MS2Query searches for both exact matches and analogues in a reference library. First, potential candidates are selected based on MS2Deepscore, followed by reranking the spectra by using a random forest model.

**Figure 2:**
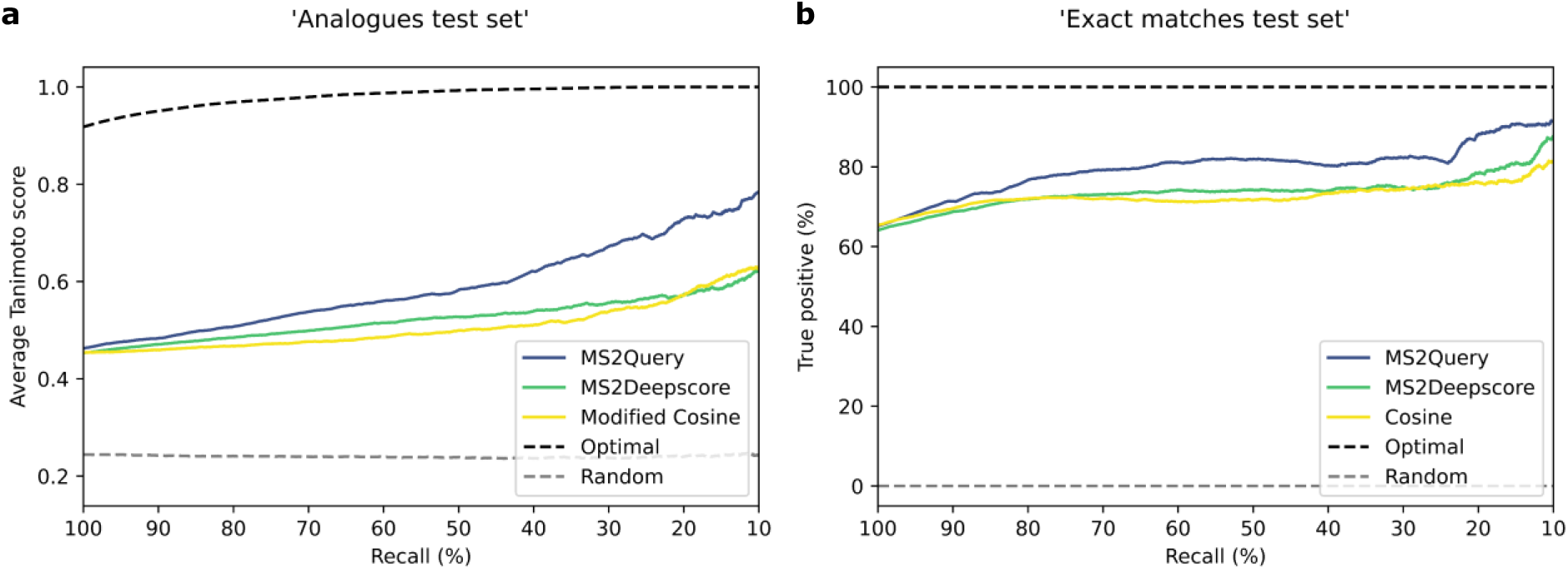
MS2Query is more accurate for finding analogues and exact matches than using MS2Deepscore or (modified) cosine score. The threshold for MS2Query, MS2Deepscore and (modified) cosine is varied, resulting in different recalls. **a**: The ‘analogues test set’ is used with spectra that have no exact match in the library, therefore the best possible match is always an analogue. For MS2Deepscore and modified cosine score, library spectra are first filtered on a mass difference of 100 Da. The relationship between recall and accuracy is plotted. For each threshold the accuracy is measured by taking the average over the Tanimoto scores (chemical similarity) between the correct molecular structure and the predicted analogues. **b**: The ‘exact match test set’ of 3,000 spectra is used, all these test spectra have at least 1 exact structural match in the reference library. For MS2Deepscore and modified cosine score, library spectra are first filtered on a mass difference of 0.25 Da, while MS2Query does not use any pre-filtering on mass difference, and uses the exact same settings as for the analogue search. The percentage of true positives is given, a match is marked as true positive if the first 14 characters of the InChiKeys are identical.

To determine the performance for finding an exact match, the percentage of predictions that is an exact match for the test spectra is calculated for the ‘exact match test set’ (Figure 2a). For MS2Deepscore and cosine score the preselection on precursor m/z difference was set to 0.25 Da, while for MS2Query no pre-filtering on mass difference was used, since MS2Query used the exact same settings and model as for the analogue search. Figure 2b shows that MS2Query performs better at finding exact matches compared to search methods relying on MS2Deepscore or the cosine score.

### Increased performance for larger metabolites

Many tools for MS^2^ spectrum annotation do not perform equally well for low and high query masses^10, 23, 29^. For MS2Query the performance for different masses is tested by splitting the ‘analogues test set’ with spectra without an exact match in the library into three mass ranges; 0-300 Da, 300-600 Da and > 600 Da. Figure 3 displays the Tanimoto score distributions of the suggested analogues for these three mass ranges. This analysis reveals that MS2Query performs best for large metabolites (>600 Da) where it detected analogues with an average Tanimoto score of 0.85 and has a recall of 63%. A better performance for larger metabolites can also be observed when using MS2Deepscore or modified cosine score, see Figure S1 and S2. However, in comparison to MS2Deepscore and modified cosine score, MS2Query was able to filter out more bad analogues for lower masses.

**Figure 3:**
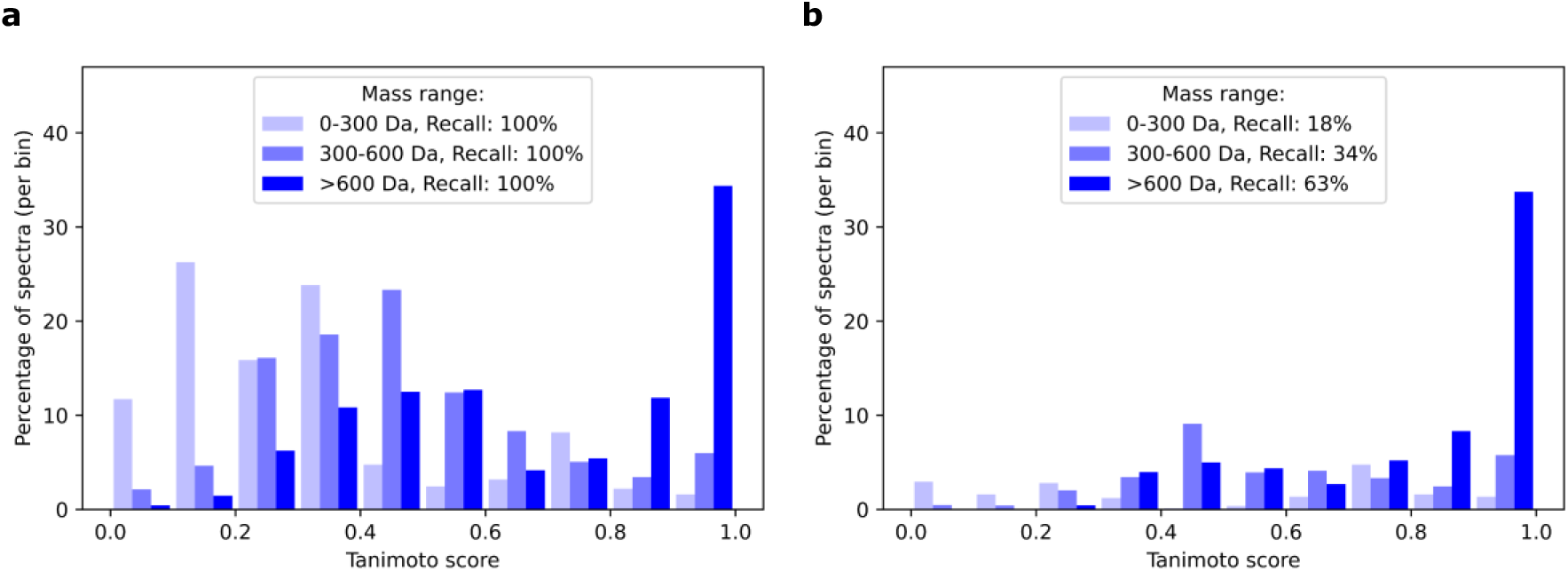
The performance of MS2Query is substantially better for larger metabolites than smaller metabolites. The ‘analogues test set’ without exact library matches is split into 3 mass bins; 0-300 Da, 300-600 Da and 600-2,000 Da. The highest scoring library spectrum is selected and the Tanimoto score is calculated between the predicted library molecule and the correct molecular structure **a**: Performance of MS2Query without using a minimal threshold for the MS2Query score. **b**: A minimal threshold for MS2Query of 0.633 was used, resulting in a total recall of 35% across the three mass bins.

### Case studies on experimental datasets of complex metabolite mixtures

MS2Query was run on four case studies, to demonstrate that MS2Query also performs well on newly generated experimental data. A urine sample, two blood plasma samples, and an anammox bacterial sample set were analysed using MS2Query and GNPS analogue search. The results of the case studies were manually validated and partially confirmed by in-house reference standards. Though informative, we would like to stress that a fair comparison of the performance in these case studies is challenging, since often no ground truth can be found for all spectra and judging whether two chemical structures are analogues remains to some extent subjective. For all case studies, the detailed results can be found in the supplementary information. Below we highlight some of the results of four case studies to illustrate that MS2Query is able to predict useful exact matches and analogues for newly generated data.

Figure 4.1a shows the number of spectra for which MS2Query predicted a match (recall) for the four case studies. The recall for the four case studies is highly variable, but on average, the case studies do not have a clear higher or lower recall compared to the benchmarking test set used. Figure 4.1b shows that the ratio between the number of predicted analogues (mass difference >1 Da) and predicted exact matches (mass difference <1 Da) differs between the case studies. Manual validation shows that most predictions by MS2Query were analogues or exact matches that matched with prior biochemical knowledge on the sample (Figure 4.2). This confirms that MS2Query is able to generate relevant predictions for newly generated experimental data.

**Figure 4:**
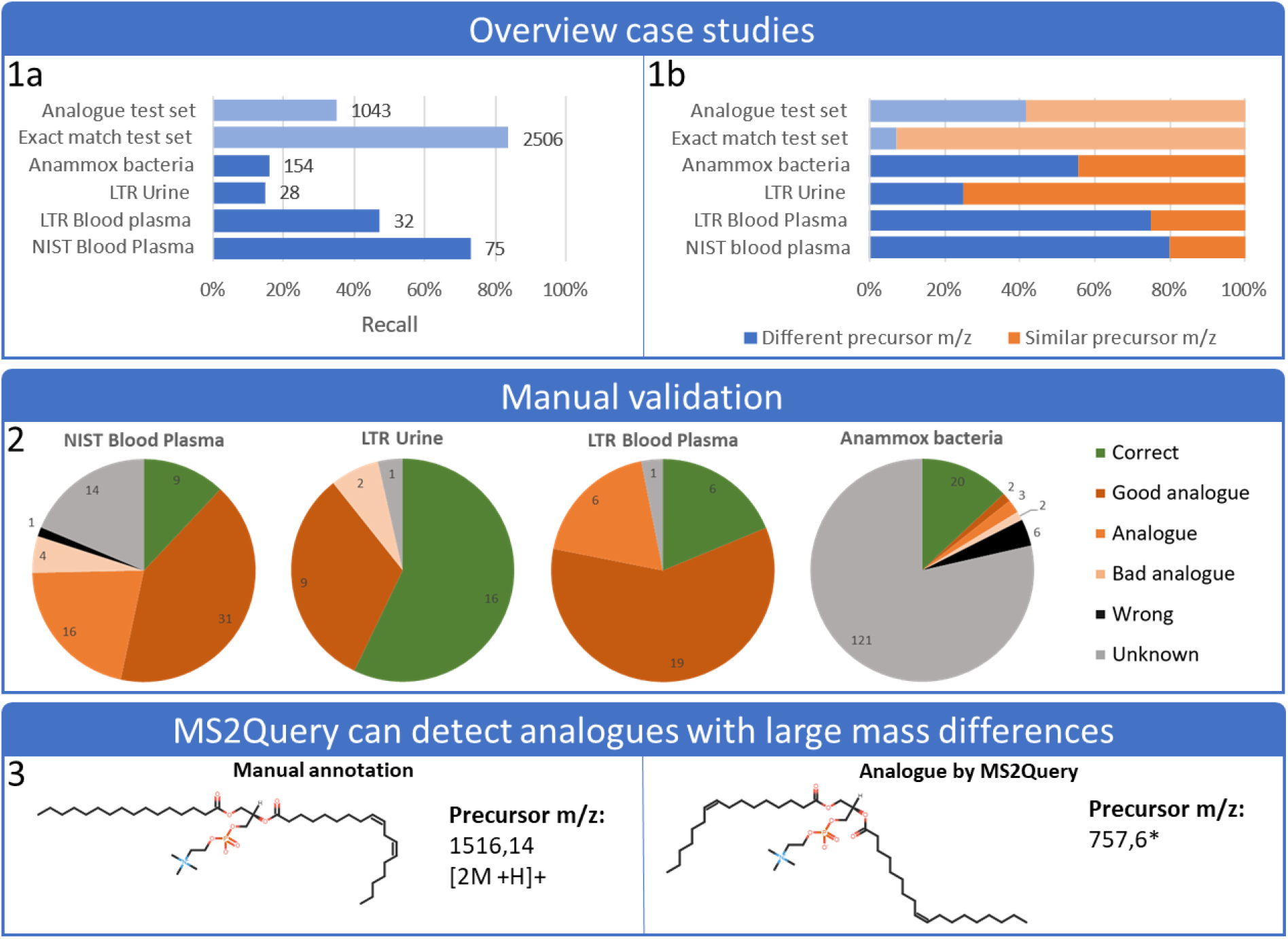
Highlight of the results of the case studies. A minimal threshold of 0.633 for the random forest score was used to determine if an analogue was selected. The threshold of 0.633 was selected, since this resulted in a recall of 35% for the “analogue test set”. **1a**: The variation of recall across case studies using the same settings. **1b**: The percentage of query spectra with a predicted analogue (precursor mz >1 Da) is compared to the percentage of spectra with an exact match predicted (precursor mz <1 Da) **2**: Results were manually validated based on the retention time MS1 mass and MS2 spectra, by comparing to online libraries or in-house reference standards. These reference standards were used to judge the quality of the predicted analogues. In the supplementary information more details about the validation can be found. For the anammox bacteria sample set, tentative validation was attempted for 50 features. **3**: MS2Query can detect analogues with large mass differences, this makes it possible to also detect analogues that have large mass differences (such as dimers for instance). The correct annotation was PC(16:0/18:2) and the predicted analogue was PC(16:1/18:1). *The precursor m/z of the predicted analogue was incorrectly stored in the GNPS library, since the parent mass of 757,6 Da was stored in the library instead of the precursor m/z of 758,6 Da.

The NIST plasma sample analysed by lipid profiling assay in positive mode contained 139 spectra for which MS2Query predicted 75 matches. Since this blood plasma sample was analysed by lipid profiling assay it was expected to contain mainly lipids. In line with this expectation, MS2Query predicted 72 matches (out of 75) to be lipids. This shows that MS2Query is able to reliably find analogues which consistently match the correct compound class.

## Discussion

Structural elucidation based on mass spectrometry fragmentation data remains hampered by a limited number of reference mass spectra in spectral libraries. Only a fraction of the mass spectra in experimental data can therefore be annotated. Many different approaches target this structural annotation problem, for instance fragmentation tree based methods^10-12^, or approaches generating in silico spectra based on structural libraries^13, 14^. Even though these are promising approaches, the problem of automatically assigning structures to mass spectra remains unsolved. Searching for so-called analogues is an attractive alternative to exact library matching. Analogues are library molecules which are not exact matches, but are structurally very similar. Analogues can be used as a starting point for complete annotation, to select metabolites of interest, or for direct biological interpretation. A benefit of searching for analogues compared to compound class prediction is that analogues make interpreting more flexible. The choice is not limited to specific compound classes, but can be extended to specific side groups for metabolites of interest, involvement in certain pathways, or relatedness to specific drugs or contaminants. Furthermore, searching for analogues can potentially help in efficiently increasing the chemical diversity of public libraries. If an analogue search does not return any matches, this metabolite is likely to be unrelated to known metabolites. Prioritizing such metabolites for structural identification by NMR spectroscopy would be an efficient way to increase the chemical diversity of public libraries. Here we introduce MS2Query a tool that is able to search a large mass spectral library both for exact matches and analogues. Based on the performed benchmarking, we expect that searching for analogues in currently publicly available mass spectral libraries, MS2Query will typically result in useful analogues for about one third of all compounds of a complex sample. The precise fraction, however, will vary depending on the exact composition and origin of a sample and the similarity of its molecules with those in mass spectral libraries.

Comparison with (modified) cosine score and MS2Deepscore shows that MS2Query performs better both at finding exact matches as well as finding analogues for positive mode MS^2^ spectra. Using a modified cosine score is a common approach for doing an analogue search, for instance implemented on GNPS^4^ and MASST^9^. Even though we demonstrate that MS2Query is able to rapidly provide reliable analogues for unknown substances, there is still room for improvement. The current version was trained using available data from GNPS^16^. While a very valuable resource, we do expect that our models will notably improve when our library is built from larger and chemically more diverse datasets. The dependency on enough and diverse training data is clearly visible when using MS2Query on negative mode mass spectra. For negative mode mass spectra, MS2Query performed less well (supplementary information S7), which is probably due to the lower number of publicly available mass spectra in negative mode. Nevertheless, MS2Query currently represents a substantial step forward in reliability, thereby creating new opportunities to use analogues to get more reliable insights into unknown mass spectra.

Our four case studies show how the number of spectra for which MS2Query predicts a match (recall) varies from 15 to 75% with the same settings (Figure 4). The observed variation can be due to differences in the quality of the acquired spectra, the masses of the metabolites, or the differing similarity between the metabolites in the sample and the metabolites in the reference libraries. This, in combination with the challenges of manually validating results, makes it hard to objectively judge if MS2Query performs similarly on newly generated data, compared to the benchmarking test set. Nonetheless, the case studies show that MS2Query is able to generate useful results for newly generated experimental data and that it can contribute to new biochemical insights based on previously unconnected analogues.

MS2Query performs particularly well at predicting analogues for molecules larger than 600 Da (Figure 3). A likely explanation why analogue searching is more accurate for larger metabolites, is that larger metabolites will often produce a higher number of characteristic fragments. In addition, small chemical changes will have a more severe impact on fingerprint based chemical similarity for smaller molecules, since those changes can quickly alter a large fraction of a fingerprint. In practice, the observed high analogue similarity for larger molecules is very promising, since it is complementary to currently existing methods relying on fragmentation tree-based approaches. Fragmentation tree-based methods perform well for smaller metabolites <500 Da, but perform less well for larger metabolites, both in terms of computational time and reliability^10, 29, 30^. This shows the potential for combining the two approaches and using the best of both for optimal performance.

Since a preselection on MS2Deepscore is the start of our method, the improved performance of MS2Query compared to MS2Deepscore shows the added value of using the five features and the random forest for re-ranking the library spectra. Additional analysis of the feature importance indicates that each of the five features used contain relevant information for correctly ranking candidate structures (supplementary Table 1 and 2). Besides the five used features, multiple other features were tested as well, for instance the cosine and modified cosine score. These other features were not selected, since they did not improve the performance of the model. Details about the other features that were tested can be found in the supplementary information S2.

MS2Query is available as an easily installable python package, which is stable and well-tested. The model as well as the library mass spectra used are available on Zenodo. MS2Query is fully automatic and was designed with the end-user in mind. For example, it outputs a CSV file with all relevant information about the found matches for the query spectra. For each found analogue it also returns the chemical compound classes based on ClassyFire^31^ annotations, to make it easier to interpret the results. MS2Query is optimized for speed and working memory usage, which makes it possible to run MS2Query on a normal laptop on 1,000 spectra within 13 minutes against a reference library of 302 514 spectra, without doing any preselection on precursor m/z difference. The scalability of MS2Query is an encouraging step toward higher-throughput large-scale untargeted metabolomics workflows, thereby creating the opportunity to develop entirely novel large-scale full sample comparisons.

MS2Query is a tool able to search for both analogues and exact matches in large spectral libraries. The tool is scalable and has an improved accuracy compared to conventional methods, this makes MS2Query very suitable for high throughput analysis. The good performance of MS2Query for larger metabolites offers a lot of new opportunities for further increasing the metabolite annotation rate of complex metabolite mixtures, in particular for natural product relevant mass ranges.

## Online Methods

### Workflow MS2Query

MS2Query builds on the improvements of two machine learning-based methods, developed to predict chemical similarity from MS^2^ mass spectral pairs; Spec2Vec^23^ and MS2Deepscore^25^. These methods perform especially well at predicting chemical similarity for molecules that are similar, but are chemically not exact matches. This makes these scores very suitable for an analogue search.

The workflow for running MS2Query first uses MS2Deepscore to calculate spectral similarity scores between all library spectra and a query spectrum. The top 2,000 spectra with the highest MS2Deepscore are selected. To optimally rank these 2,000 spectra, MS2Query calculates 5 features which are combined by a random forest model. The prediction of the random forest model is used to rank the 2,000 preselected library spectra (See Figure 1). As input for the random forest model, MS2Query uses 5 different features, calculated between the query spectrum and each of the 2,000 preselected library spectra. The features are 1. Spec2Vec similarity, 2. query precursor m/z, 3. precursor m/z difference, 4. an average MS2Deepscore over 10 chemically similar library molecules, and 5. the average Tanimoto score for these 10 chemically most similar library molecules.

The **Average MS2Deepscore** of multiple library molecules (feature 4), builds on the following principle. For two library molecules that are chemically very similar, it is expected that if one of these library molecules is a good analogue to your query spectra, the other is a good analogue as well. For this reason it is expected that for a good analogue the MS2Deepscore between such a chemically similar library molecule and your query spectrum is also high. This is captured in this feature by calculating the average MS2Deepscore between a query spectrum and all spectra of 10 chemical similar library molecules (Figure 5). These 10 library molecules are selected based on the known chemical structures of the spectra in the library, by selecting the library structures with the highest Tanimoto score. For each of the 10 library molecules all corresponding library spectra are selected. The MS2Deepscore between these library spectra and the query spectrum is calculated and the average per library structure is taken. As an input feature for the random forest model, the average over the MS2Deepscore for the 10 library structures is used (Feature 4). In addition, the average of the Tanimoto score between the starting library structure and the 10 library structures is used as an additional input feature (Feature 5).

**Figure 5:**
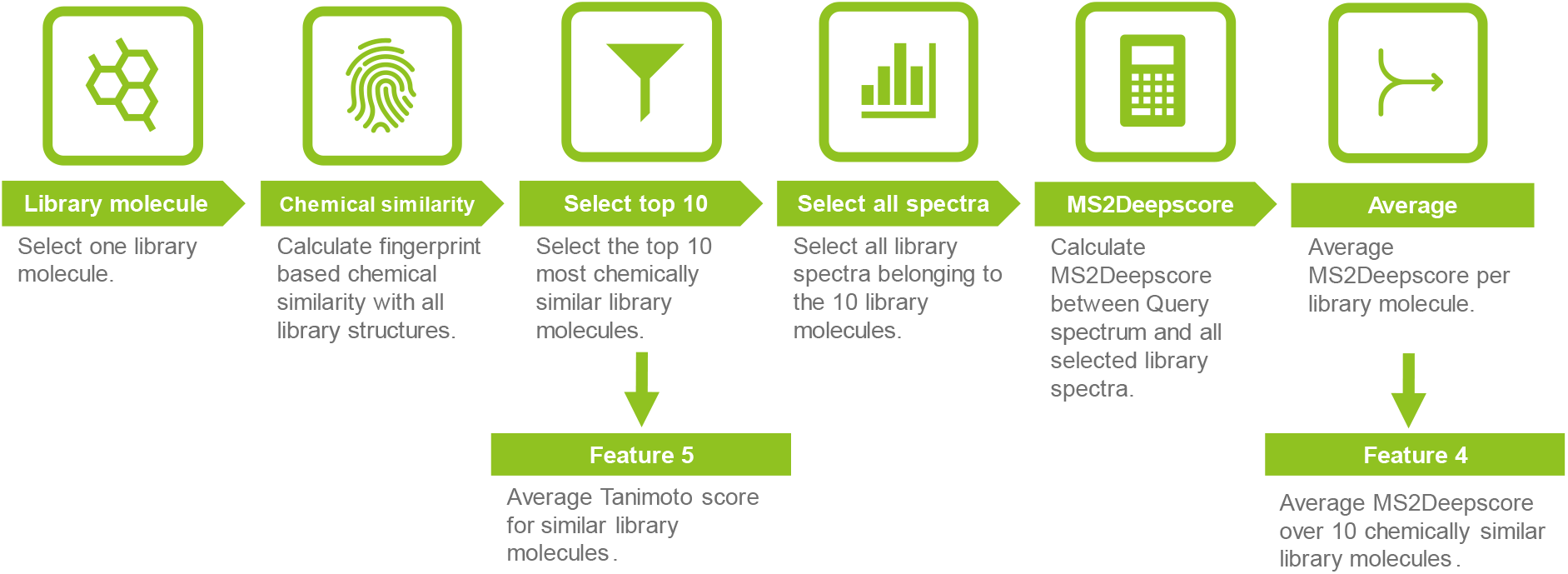
Workflow for calculating two input features of the random forest model. Feature 5 is the Average Tanimoto score for similar library molecules and feature 4 is the average MS2Deepscore over 10 chemically similar library molecules.

### Tanimoto scores as structural similarity label

First, an rdkit^28^ daylight fingerprint (2,048 bits) is generated from the InChI for each unique 14-character InChIKey in the library. If multiple spectra with the same InChIKey exist in the dataset, the most frequently occurring InChI was selected and used for all spectra with the corresponding InChIKey. A Tanimoto score^27^ was calculated between the molecular fingerprints for each pair of InChIKeys. The Tanimoto score is used as an indication for structural similarity of that pair. These Tanimoto scores are used as labels for training MS2Deepscore and MS2Query and for selecting chemically similar library molecules to calculate an average of the MS2Deepscore of multiple chemically similar library spectra.

### Data cleaning

For training and testing of MS2Query, we used data from GNPS. The GNPS dataset used was downloaded from GNPS (https://gnps-external.ucsd.edu/gnpslibrary/ALL_GNPS) on the 15^th^ of November 2021, 20:00 CET. The dataset was first cleaned using matchms^26^. The metadata was cleaned to get a uniform format and to remove or correct misplaced metadata. The intensities of the mass fragmentation peaks are normalised. Peaks above 1,000 Da were removed and peaks with an intensity of less than 0,1 % of the highest peak were removed. For spectra with more than 500 peaks, the peaks with the lowest intensities were removed. Spectra with less than 3 peaks were completely removed from the library. Some spectra in the GNPS library do not have an InChIKey stored. A method from matchms extras was used to add missing InChIKeys by searching the compound name and molecular formula on PubChem. The number of spectra at each step can be found in Figure 6.

**Figure 6:**
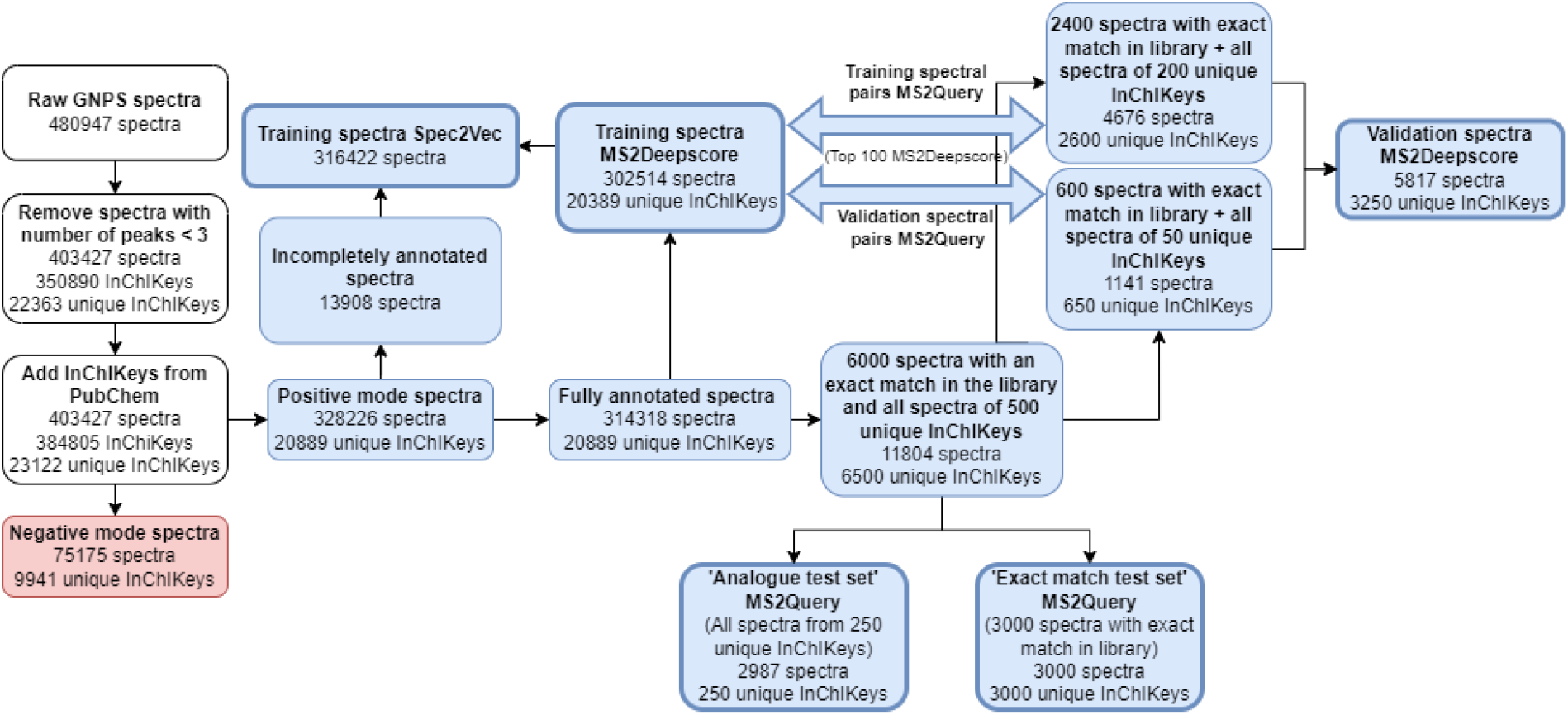
Workflow for creating datasets used for training, validation and testing of MS2Deepscore, Spec2Vec and MS2Query for spectra in positive ionization mode.

**Figure 7:**
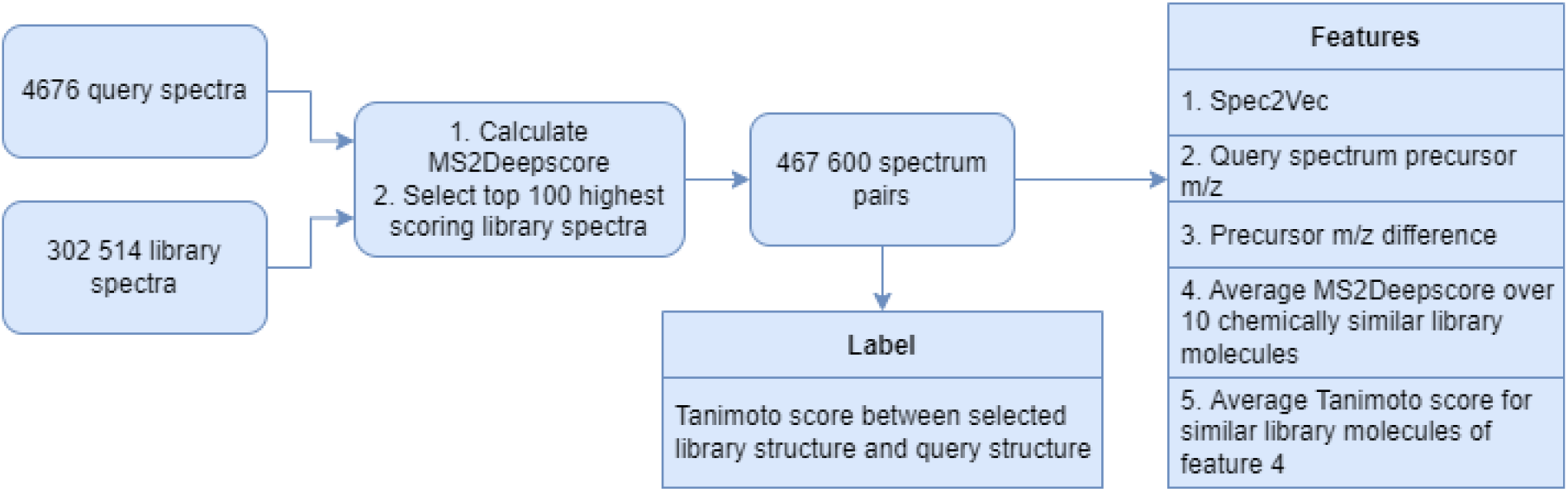
Workflow for training random forest model of MS2Query.

The spectra are split in two separate datasets for spectra obtained in positive and negative ionization mode. The processing and benchmarking of the negative mode spectra can be found in supplementary information S7. For all other benchmarking only the positive mode spectra are used. These spectra are split into a training set and two different test and validation sets that address different goals of MS2Query. For both the test and validation set, 250 random InChIKeys were selected, for which all spectra were removed from the library. In addition, 3,000 random spectra were selected that belong to an InChIKey that has more than one spectrum in the library. The two validation data sets are added together during training of MS2Deepscore and Spec2Vec.

### Training MS2Deepscore and Spec2Vec

MS2Deepscore was trained on all fully annotated spectra from the GNPS library, using the same settings as used for the MS2Deepscore publication^25^.

A spec2vec model is trained using all spectra from the GNPS library, both annotated and unannotated spectra. The model is trained in 30 epochs using binning on 2 decimals^23^.

### Training random forest model

The random forest model used by MS2Query was trained on pairs of annotated spectra using 5 different features. The model was trained to predict the Tanimoto score between the two structures of each pair. In total 467 000 spectral pairs are used as input for training the random forest model. To generate the training spectrum pairs 2,400 spectra with at least 1 library match + all library spectra of 200 InChIKeys (2,276 spectra) where removed from the library spectra. The spectrum pairs were generated by starting with one spectrum from this set and creating spectrum pairs with the 100 library spectra that have the highest scoring MS2Deepscore for this spectrum. As starting point to generate the validation spectrum pairs 600 spectra with at least 1 match + all spectra of 50 InChIKeys (541 spectra) were removed from the library. More details about the motivation for selecting the top 100 highest scoring spectra for training can be found in supplementary information S5.

The implementation of scikit-learn^32^ was used for the random forest model. The mean squared error was used as a loss function. The number of estimators was set to 250 and the max depth to 5. The implementation of scikit-learn was used to calculate the feature importance of the 5 scores used. This method is based on an impurity-based feature importance, also known as the Gini importance^33^.

Beside these 5 features, multiple other features were tested as well, for instance the cosine and modified cosine score. These other features were not selected, since these did not improve the performance of the model. Details about the other features tested and the distributions of Tanimoto scores in the training data can be found in the supplementary information S2.

### Benchmarking

MS2Query was designed to search for analogues and exact matches in one run. Since these goals are slightly different, they were both benchmarked separately. The performance for an analogue search was benchmarked by using a test set that does not have any exact match in the library, therefore the best possible match will always be an analogue. To benchmark the search for exact matches a dataset was used that always has an exact match in the library. See Figure 6 and the section Data cleaning for the exact method for creating these datasets.

### Analogue search

MS2Query, MS2Deepscore, or the modified cosine score were used to rank the reference library spectra. For MS2Deepscore and modified cosine score, library spectra were first filtered on a maximum precursor m/z difference of 100 Da. For MS2Query, spectra were not filtered on precursor m/z difference. The Tanimoto score between the predicted library molecule and the correct test molecule was calculated and used as a performance metric of each method. All three methods use a minimal threshold for the spectral similarity score to determine if a library spectrum is a good analogue. The threshold for each method was varied between 0 and 1, followed by calculating the accuracy and recall.

### Exact matches

MS2Query, MS2Deepscore, or the cosine score were used to rank the reference library spectra. For MS2Deepscore and cosine score only library spectra were considered within a mass tolerance of 0.25 Da. For MS2Query no minimum threshold for mass tolerance was used. The cosine greedy implementation of matchms^26^ was used to calculate the cosine score, a fragment mass tolerance of 0.05 Da was used. The InChIKey of the predicted molecule was compared to the InChIKey of the correct molecule. If the first 14 letters of both InChIKeys were the same, a found match was marked as correct, if the first 14 letters differ it was marked as wrong.

### Separation of mass

The test dataset for the analogue search was split into 3 mass categories. Spectra with a precursor m/z of 0-300 Da, of 300-600 Da and all spectra larger than 600 Da. The size of these 3 seperate test sets were 819, 1,684 and 481 spectra respectively. The performance of these 3 test sets was determined separately.

### Speed and memory optimization

MS2Query was optimised for speed and working memory efficiency. To make this possible, MS2Query aims to avoid repetitive, computational expensive operations. The biggest speed improvement was achieved by pre-calculating mass spectral embeddings for Spec2Vec and MS2Deepscore. MS2Deepscore and Spec2Vec both predict a chemical similarity score between two library spectra, by first calculating a multidimensional embedding followed by calculating the (mathematical) cosine similarity between these two embeddings. The library spectra are already known, therefore the embeddings for all library spectra are pre-calculated and stored. Therefore only for the query spectra the embeddings have to be computed, instead of all the library spectra.

In the first step the top 2,000 library spectra are selected that have the highest MS2Deepscore between the query spectrum and a library spectrum. To do this selection, the MS2Deepscores between a query spectrum and all MS2Deepscores are calculated. To avoid repetitive calculation of these scores, the calculated MS2Deepscores are reused to calculate the average of the MS2Deepscore of multiple chemically similar library molecules.

The precursor m/z is the only metadata entry that is required for MS2Query and which serves to calculate the mass differences. Other spectra metadata such as retention time, SMILES or compound names can be returned for results found by MS2Query. To reduce the toll on working memory, this information is stored in a SQLite library. The precursor m/z is stored in a separate SQLite library column for efficient look-up speeds. To calculate the average MS2Deepscore of multiple chemically similar library molecules, the 10 most chemically similar library molecules based on the Tanimoto score are needed. This top 10 list of most related InChIKeys is pre-calculated for every unique InChIKey in the library and stored in the SQLite library.

MS2Query contains a workflow to automatically generate all needed files, making it straightforward to recreate these files for new or different spectral libraries.

### Speed performance

The speed was tested on the 5,987 test spectra in positive mode and compared to the positive mode GNPS library containing 302.514 spectra. The test was run on a laptop; the Lenovo Thinkbook 15-IIL. This laptop has an 11th generation Intel Core i5-1135G7 and 16GB installed RAM.

### Case studies

Four case studies were performed to confirm that MS2Query performs well on newly generated experimental data. Two blood plasma samples, a urine sample and a bacterial sample set were analysed. The raw data, intermediate files and raw results can be found on https://zenodo.org/record/6811540#.YshH5GBBxPY. Here below, the analytical methods used, and the data preprocessing and processing steps are described for all case studies.

### Case study 1: NIST Human blood plasma

For this case study, the NIST 1950 Frozen Human Plasma standard reference material (SRM). The sample was subjected to reversed-phase chromotographic (RPC) assay tailored for complex lipid separation as described in Lewis, *et al*.^34^.

### Case study 2: Blood plasma Long-Term Reference

For this case study, a plasma Long-Term Reference (LTR) sample was used. This LTR is routinely integrated in profiling studies at the National Phenome Centre for study-independent monitoring of precision. To create the plasma LTR, 10 L of bulk plasma were purchased from Seralab, homogenized, and aliquoted for long term storage at –80ºC. Hydrophilic interaction liquid chromatography (HILIC) was used in this case study for the analysis of polar metabolites in a sample of plasma LTR^34^.

### Case study 3: Urine long-Term Reference

For this case study, a pooled Long-Term Reference (LTR) urine sample, maintained by the National Phenome Centre and utilized as an independent sample reference throughout all molecular profiling studies, was used. The protocol followed to generate urine LTR sample is described in detail by Lewis *et al*.^35^. Briefly, this material was created by pooling together 78 urine voids collected from healthy volunteers in one day. All samples were combined in a 20 L vessel, homogenized and aliquoted into 15 mL polypropylene conical centrifuge tubes (Corning) for long term storage at –80ºC. Samples were analyzed by reversed-phase chromatographic (RPC) assay tailored for small molecule metabolites^34^.

### Preprocessing Case study 1

NIST 1950 human plasma sample was thawed at 4°C for 2h. Subsequently, a 50 μL aliquot was taken and prepared for lipid analysis by dilution with LC-MS grade water (1:1 v/v) and addition of four parts of isopropanol (IPA) containing a mixture of lipid reference standards^34^ to one part of diluted sample for protein precipitation. Vial with the sample was mixed at 1,400 rpm for 2h at 4°C and subsequently centrifuged for 10 mins at 3,486×g at 4°C to separate the homogenous supernatant from the precipitated protein. The clear supernatant was aspirated and dispensed into LC-MS vial, then additionally centrifuged for 5 mins at 3,486×g and 4°C prior analysis. Prepared sample was injected (1μL) in the chromatographic system using full loop mode (5× overfill).

### Preprocessing Case study 2

For the case study 2, plasma LTR sample was prepared for the analysis by HILIC method in positive ionization mode. The sample was thawed at 4°C for 2h. Subsequently, a 50 μL aliquot of plasma LTR sample was diluted 1:1 with LC-MS grade water and HILIC internal standards (IS)^34^. Three parts of acetonitrile were then added to one part of diluted sample for protein precipitation. Vial with the sample was mixed at 1,400 rpm for 2h at 4°C and subsequently centrifuged for 10 mins at 3,486×g at 4°C to separate the homogenous supernatant from the precipitated protein. The clear supernatant was aspirated and dispensed into LC-MS vial, then additionally centrifuged for 5 mins at 3,486×g and 4°C prior analysis. Prepared sample was injected (2μL) in the chromatographic system using full loop mode (5× overfill).

### Preprocessing Case study 3

Preparation and analysis of urine samples are described in detail by Lewis *et al*.^35^. In brief, an aliquot of 150 μL of urine sample was diluted with 75 μL of ultrapure water and 75 μL of RPC-specific internal standards (IS) solution^34^. The sample was mixed at 850 rpm for one minute at 4°C and centrifuged for 10 mins at 3,486×g at 4°C. The supernatant was aspirated and dispensed into LC-MS vials for the analysis. Urine sample was injected (2μL) in the chromatographic system using full loop mode (5× overfill).

### UPLC-MS profiling analysis for case studies 1-3

All UPLC-MS analyses were performed on Acquity UPLC instruments, coupled to Xevo G2-S TOF mass spectrometers (Waters Corp., Manchester, UK) via a Z-spray electrospray ionization (ESI) source.

For lipid profiling, all solvents – water, acetonitrile (ACN), and IPA and mobile phase additives ammonium acetate and acetic acid were of LC-MS grade. Lipidomic profiling was conducted using a 2.1×100 mm BEH C8 column, thermostatted at 55°C. Mobile phase A consisted of a 2:1:1 mixture of water:ACN:IPA with 5mm ammonium acetate, 0.05% acetic acid, and 20 μM phosphoric acid. Mobile phase B consisted of 1:1 ACN:IPA with 5 mM ammonium acetate, 0.05% acetic acid. The initial conditions were 99:1 A:B at a flow rate of 0.6 mL/min. The details of gradient elution program are shown in the protocols associated with Lewis *et al*.^34^.

The HILIC chromatographic retention and separation of polar molecules was conducted using a 2.1 × 150 mm Acquity BEH HILIC column thermostatted at 40 °C. 20 mM ammonium formate in water with 0.1% formic acid was used as mobile phase A and ACN with 0.1% formic acid as mobile phase B. The initial conditions were 5:95 A:B at a flow rate of 0.6 mL/min. The details of gradient elution program are shown in the protocols associated with Lewis *et al*.^34^.

For urine profiling, water and ACN supplemented with 0.1% formic acid of LC-MS grade were used as mobile phases A and B. A 2.1 × 150 mm HSS T3 column thermostatted at 45 °C was used with a mobile phase flow rate of 0.6 mL/min. The details of gradient elution program are shown in the protocols associated with Lewis *et al*.^34^.

The analysis of blood plasma and urine reference samples in presented case studies were performed in positive ionization mode. The mass spectrometry parameters were set as follows: capillary voltage 2 kV for lipid profiling and 1.5 kV for urine profiling, sample cone voltage 25 V for lipid profiling and 20 V for urine profiling, source temperature 120°C, desolvation temperature 600°C, desolvation gas flow 1,000 L/h, and cone gas flow 150 L/h. Data were collected in centroid mode with a scan range of 50-2,000 *m/z* and 50-1,200 *m/z* for lipid and urine profiling, respectively, and a scan time of 0.1 s. For mass accuracy, LockSpray mass correction was performed using a 600 pg/μL leucine enkephalin solution (*m/z* 556.2771 in ESI+) in 1:1 water:ACN solution at a flow rate of 15μL/min. Lockmass scans were collected every 60 s and averaged over 4 scans. The mass spectrometer was operating in Fast DDA mode. The intensity threshold of precursor ion was set to 100 K to trigger MS^2^ fragmentation that was performed in centroid mode with a scan range of 50-2,000 m/z and a scan time of 0.25 s. MS^2^ was switched back to MS survey function after 2 s acquisition. Deisotoped peak selection option was enabled. The collision energy was set to the ramp of 15–30 eV and 30-60 eV for MS^2^ acquisition of low and high mass ions, respectively. Ten iterative DDA acquisitions were performed using DDA auto exclude program, which allows ions selected as precursors in previous injections are removed from the list in the following injections.

### Case study 4: Anammox bacteria

For the fourth case study extracts of three strains of anammox bacteria were used. *Kuenenia stuttgartiensis* MBR1 was cultivated in a 12 liter single cell membrane bioreactor (MBR) as previously described by Kartal *et al*.^36^. *Brocadia fulgida* was cultivated as previously described by Kartal *et al*.^*37*^ with some adjustments: the working volume was 6 liter and the bacteria were kept in a single cell membrane bioreactor. *Scalindua* was cultivated in a 5.5 liter sequencing batch reactor at room temperature as described earlier by van der Vossenberg *et al*.^38^. Samples (30 mL) were taken from each reactor in triplicate and kept on ice. After centrifugation at 3,000 x g, at 4 °C for 5 minutes, the cell pellets were lysed in ice-cold acetonitrile:methanol:water (2:2:1; v:v:v). The samples were snap frozen in liquid nitrogen and stored at -70°C until further use. To remove precipitated proteins and extracellular matrix, samples were centrifuged again at 20,238 x g, at room temperature for 5 minutes. Subsequently, samples were subjected to LC-MS analysis as described previously by Jansen *et al*.^*39*^ with several adaptations. The samples were injected onto a Diamond Hydride Type C column and separated using a gradient of acetonitrile and water (both with 0.2% formic acid) on an Agilent 1290 II LC system coupled to an Agilent Accurate Mass 6546 Quadrupole Time of Flight (Q-TOF) instrument operated in the positive ionization mode and a scan range of 50-1,200 m/z. For data dependent acquisition of MS2 spectra, automated selection of maximum 4 precursor ions (> m/z 100) per cycle with an exclusion window of 2 minutes after a single spectrum, and an absolute threshold of 1,000 counts with a mass error tolerance of 20 ppm was used. The scan speed was varied based on precursor abundance with a target of 50,000 counts. Common background ions were excluded, the isolation width was set to narrow (∼1.3 m/z), and the collision energy was set to 20 V. Data collection was performed using Agilent Masshunter software 10.0 (Agilent Technologies).

### Data processing

In the case of case studies 1-3 the spectra were uploaded on GNPS to run MSCluster^40^ to create consensus spectra. These consensus spectra were taken as input for MS2Query. The data files for case study 4 were first converted to mzML format using Proteowizard (Chambers et al., 2012). Next, LC-MS features were picked using XCMS3^41^ (https://github.com/sneumann/xcms), using the findChromPeaks function. The resulting MS2 spectral MGF file was used to run MS2Query.

Analogues and exact matches with a MS2Query score above 0.633 (corresponding to 35% recall for the “analogues test set” during benchmarking) were considered for all case studies. In addition, an analogue search on the GNPS platform^16^ for case studies 1-3 was performed and FBMN for case study 4 was performed. More information about this can be found in the supplementary information S6.

### Manual validation

To validate the MS2Query matches for case study 1-3, metabolites with MS^2^ were manually annotated to confidence level 1-3 according to the Metabolomics Standards Initiative^42^ by matching fragmentation spectra to reference data from an in-house standards database and online databases LIPID MAPS^43^, HMDB^44^, and GNPS^16^. In the case of case study 4, annotations were checked based on a combination of biological knowledge and matching of MS1 mass and retention time to reference standards. Judgement of the analogue quality was done manually. Lipids where the lipid type (e.g. PC or SM) was correctly predicted and the chain lengths were similar, were marked as a good analogue. Correctly predicted lipids, but wrong lipid types were marked as analogue. The detailed manual annotations and judgements for all spectra can be found as an excel file in the supplementary information for all case studies.

## Supporting information

Results and manual validation case studies

## Acknowledgements

The authors would like to thank Marynka Ulaszewska, Lapo Renai and Huali Xie for sharing experimental data to test the first versions of MS2Query. The authors would also like to thank colleagues from the eScience Center and Marnix Medema from the WU Bioinformatics Group for useful discussions related to MS2Query.

## Code and data availability

MS2Query is available as an easily installable Python library running on Python 3.7 and 3.8. Source code and installation instructions can be found on Github (https://github.com/iomega/ms2query). The presented results were obtained using version 0.3.2. The models and spectra files used can be downloaded from https://zenodo.org/record/6124553#.YrlpREZBxPY. A detailed overview of the code used for all methods can be found in the notebooks and scripts in the Github folder https://github.com/iomega/ms2query/tree/main/notebooks/GNPS_15_12_2021.

The mass spectrometry data for case study 4 were deposited in the MassIVE repository (accession number MSV000089648).

