## Supplementary information for "MS2Query: Reliable and Scalable MS^2^ Mass Spectral-based Analogue Search"

### S1. Feature importance

The resulting mean squared error (MSE) for the MS2Query random forest model predicting Tanimoto scores is 0.0282 for the training data and 0.0255 for the validation data. The feature importance for the different features is given in Table S1. This method is based on an impurity-based feature importance, also known as the Gini importance[1]. As an alternative method for assessing the impact the features have on the model performance, random forest models were trained lacking one of the features, to determine the difference in performance for these models (Table S2). Replacing the average MS2Deepscore of multiple library structures with the MS2Deepscore between 1 library spectrum and the query spectrum increased the MSE from 0.0255 to 0.0337 for the validation data set. This clearly demonstrates that using the average of multiple similar library spectra significantly improves the performance of the random forest model. Removing any of the other features also decreased the performance of the model.

*Table S1: Feature importance for the 5 features used by the random forest.*

| **Feature** | **Feature Importance** |
|---|---|
| Average of MS2Deepscore of multiple similar library spectra | 0.62 |
| Precursor m/z difference | 0.18 |
| Query spectrum m/z | 0.13 |
| Spec2Vec score | 0.05 |
| Average Tanimoto score for similar library spectra | 0.02 |

*Table S2: The effect on the MSE, when the random forest model is trained without one of the features.*

| **Removed feature** | **Training MSE** | **Validation MSE** |
|---|---|---|
| No feature removed | 0.0282 | 0.0255 |
| Average MS2Deepscore of multiple library structures | 0.0339 | 0.0331 |
| Precursor m/z difference | 0.0311 | 0.0283 |
| Query precursor m/z | 0.0300 | 0.0265 |
| Spec2Vec | 0.0290 | 0.0272 |
| Average Tanimoto score of similar library structures | 0.0283 | 0.0259 |

#### S1.1 Rationale behind Average MS2Deepscore of Multiple library molecules

The feature that has the biggest impact on the increased performance is the *Average MS2Deepscore of multiple library molecules.* This feature builds on the following principle: for two library molecules that are chemically very similar, it is expected that if one of these library molecules is a good analogue to your query spectra, the other is a good analogue as well. For this reason it is expected that the MS2Deepscore between a chemically similar library molecule and your query spectrum is also high, for a good analogue. MS2Query is the first mass spectral library searching method that uses this principle to re-rank candidate molecules.

#### S1.2 Rationale behind precursor m/z difference

Another newly introduced feature of MS2Query is using precursor m/z difference as an input feature for the random forest model. Current implementations of an analogue search often start with a preselection on precursor m/z followed by selecting spectra above a predefined threshold for the used similarity score[2]. A predefined precursor m/z threshold does not differentiate between precursor m/z differences as long as the mass difference falls within the set threshold. However, certain specific mass differences will be more likely than others. By using the precursor m/z difference as an input feature, the random forest can learn specific mass differences that are more likely to correspond to a good analogue or exact match. This approach has similarities to the analogue search implementation by Stephen E. Stein and colleagues: in their approach they limit the analogues to a predefined list of precursor m/z differences of commonly observed losses[3]. This is an interesting approach; however, this limits the analogues only to known losses and does not allow for analogues that have a substructure substituted by another substructure. Our random forest model is more flexible and can learn the mass differences that are an indication of a good analogue, even if these are not known mass differences.

### S2. Testing additional features for MS2Query random forest model

Five features were used as input for the current random forest model. However, multiple other features and variations of the current features were tested to select the features that contain information to predict chemical similarity. The performance of multiple models with different feature sets are compared. The feature importance is calculated for each model and used as a first guide for selecting important or relevant features, followed by training a new model with the selected features, to make sure the performance does increase. The performance of a model is measured by the mean squared error for the training and validation dataset. The code used to calculate these other features can be found in the branch https://github.com/iomega/ms2query/tree/add_cosine_to_features on the MS2Query Github repository, the notebooks for the generation of the random forest models can be found on https://github.com/iomega/ms2query/blob/add_cosine_to_features/notebooks/Analysis_with_dataset_GNPS_15_12_2021/test_features_for_RF/test_random_forest_with_49_features.ipynb. Below, alternative possible MS2Query input features are discussed in detail.

#### S2.1 (Modified) cosine scores

Both cosine scores and modified cosine scores were added as features for training random forest models to explore if they would notably contribute to the model predictions. However, the feature importance was always 0 for these scores, indicating that the model does not use the (modified) cosine score for the prediction at the set tree depth of 5. This suggests that the (modified) cosine score does not provide additional predictive power, when MS2Deepscore and Spec2Vec are part of the model as well. Therefore, it was decided to not incorporate the (modified) cosine score in the workflow of MS2Query.

#### S2.2 Weighting of the average of multiple library structures

The multiple library spectra were selected from the 10 library structures that are chemically most similar to the structure of interest, based on the Tanimoto scores. For all spectra belonging to these 10 selected library spectra we compute the MS2Deepscore towards the query spectrum. To find a good, reliable method for taking an average and for weighting each of these selected spectra, we tried different approaches and selected the method with the best performance. To assess which method works best the MSE was compared for two models having different approaches for this feature, while the other 4 features stayed constant (precursor_mz_difference, query_precursor_mz, spec2vec_score, and MS2Deepscore).

There are two approaches that we assessed for calculating the average of the calculated MS2Deepscores. The first method just takes the average over all selected spectra regardless of which library structure it belongs to and the second method first calculates the average MS2Deepscore for each selected library structure, followed by taking the average of the 10 average scores for each library structure. Since for one InChIKey the number of library spectra differs, the two methods result in a different way of weighting the spectra.

The model using the feature that first calculates the average MS2Deepscore for each selected library structure, followed by taking the average of the 10 average scores for each library structure performed the best. This method had an MSE of 0.0283 for the training set and 0.0257 for the validation set. The model using the feature where the average is just taken over all spectra had an MSE of 0.0301 for the training data and an MSE for the validation data of 0.0286. Therefore, the first method of selecting the average was selected.

#### S2.3 Weighting based on Tanimoto score

In the comparison done in S2.2 each library structure is weighted equally. In practice, however, we see variations in how chemically similar the 10 closest structures are (all measured using Tanimoto). To explore if this has any notable negative impact on our model performance, we tested a weighted average. This should give more importance to library molecules that are more similar. We hence weighted each score by the Tanimoto score and further also tested different powers of this weighting. A higher power will result in more extreme weight to spectra of more similar library structures and a low power will have less effect.

As weight, the Tanimoto score to the power of 1-3 was tested. In Table S3 the performance of the different models are shown. This shows that using weighting of the different structures based on the Tanimoto score did not improve the overall performance of the model. Therefore, no weighting is used for calculating the feature using multiple library structures.

*Table S3: Training and validation MSE for different weighting methods for the average MS2Deepscore of multiple library structures.*

| Method of weighting | Training MSE | Validation MSE |
|---|---|---|
| No weighting | 0.0283 | 0.0257 |
| Tanimoto score | 0.0284 | 0.0260 |
| Tanimoto score$^2$ | 0.0286 | 0.0262 |
| Tanimoto score$^3$ | 0.0290 | 0.0265 |

#### S2.4 Mass spectrometer instrument type

The mass spectra in the GNPS library are measured on a wide variety of mass spectra instruments. We tested if we could use this information to improve the performance of the random forest model, since it is conceivable that some features are influenced by the instrument type or by whether or not the two spectra of interest were measured on different instruments. To this end we used metadata of spectra to classify the query and library spectra as a mass spectrometer using "ToF", "Quadrupole", "Ion Trap", or "Orbitrap" setup. Instruments that were not given or used a system that could not be classified as one of the 4 mentioned techniques were not classified. For both the query spectrum and the library spectrum a binary feature is created for each of these mass spectrum instrument types. Thus, this resulted in 4 features to indicate the type of the query spectrum and 4 features to indicate the type of the library spectrum: a total of 8 features. The value was set to 1 if it was measured on this instrument type and 0 if it was not measured on this instrument type. If the instrument type could not be classified, all features were set to 0.

The feature importance of training a random forest model including these features always resulted in a feature importance of 0. This shows that with this setup, the random forest model was not able to use this information to increase the prediction quality of MS2Query. Therefore, this feature was left out in the current MS2Query workflow.

### S3. Performance of an analogue search for different mass ranges

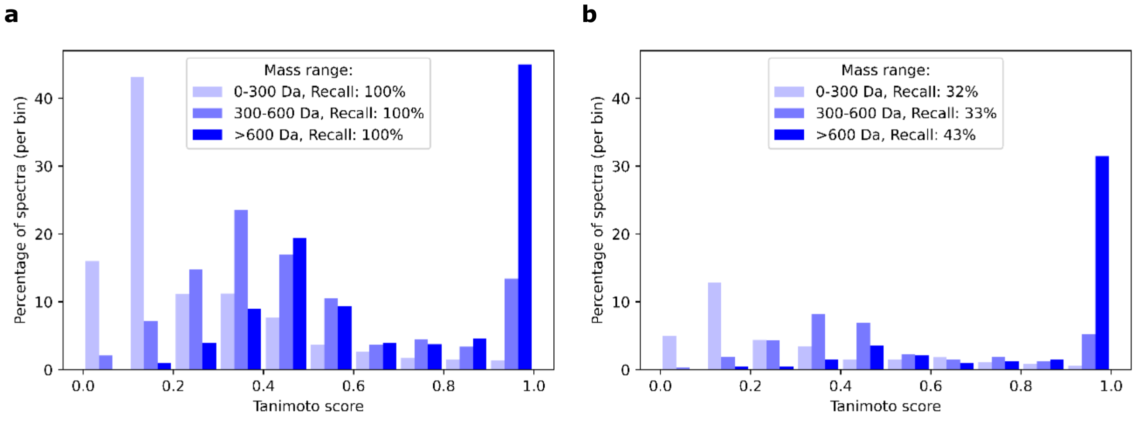


*Figure S1: Performance of Analogue search using modified cosine score and a maximum precursor m/z difference of 100 Da. The test set without exact library matches was split into 3 mass bins; 0-300 Da, 300-600 Da and >600 Da. The highest scoring library spectrum is selected and the Tanimoto score is calculated between the predicted library molecule and the actual Molecule.* ***a****: Performance of analogue search using modified cosine without using a minimal threshold for the modified cosine score.* ***b****: A minimal threshold for modified cosine score of 0.0988 was used, resulting in a total recall of 35%.*

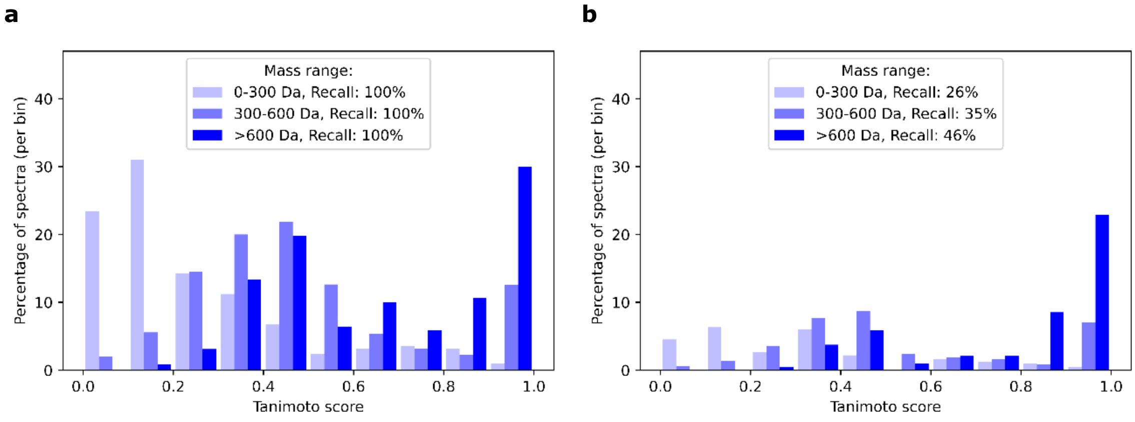


*Figure S2: Performance of Analogue search using MS2Deepscore and a maximum precursor m/z difference of 100 Da. The test set without exact library matches was split into 3 mass bins; 0-300 Da, 300-600 Da and >600 Da. The highest scoring library spectrum is selected and the Tanimoto score is calculated between the predicted library molecule and the actual Molecule.* ***a****: Performance of analogue search using MS2Deepscore without using a minimal threshold for MS2Deepscore.* ***b****: A minimal threshold for MS2Deepscore score of 0.9664 was used, resulting in a total recall of 35%.*

### S4. Detailed performance comparison and insights into analogue test set

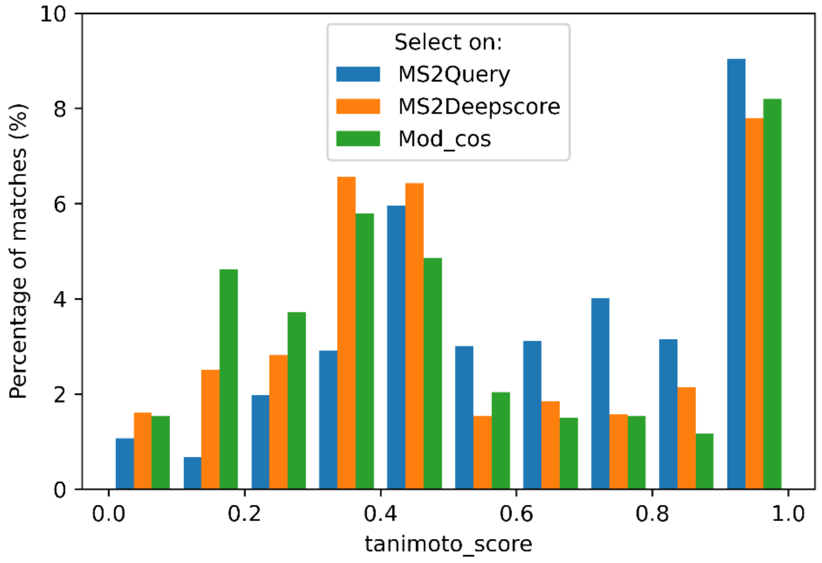


*Figure S3: The distribution of Tanimoto scores between the correct match and the best found match. The "Analogues test set" is used with spectra that have no exact match in the library, therefore the best possible match is always an analogue. The minimal threshold for MS2Query, MS2Deepscore and Modified cosine is set to result in a recall of 35% for this test set. A histogram is plotted to show the number of spectra in each subset.*

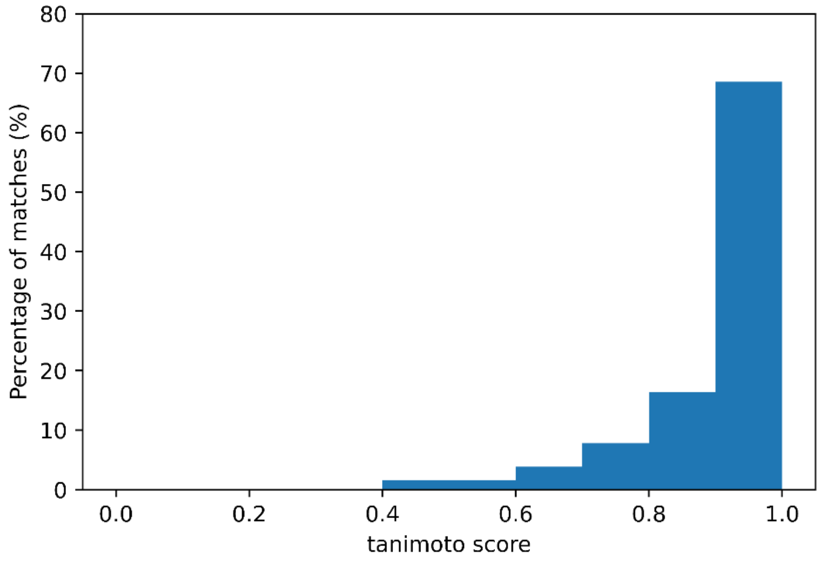


*Figure S4: Distribution of Tanimoto scores between the true structure and the best possible match in the reference library for the "Analogues test set".*

#### S5. Justification for selecting training data

Pairs of spectra were used for the training data of the random forest. If the spectra pairs would have been picked at random, the resulting Tanimoto scores would be very low on average[4]. To prevent overfitting to low Tanimoto scores, it is important that a more equal distribution of Tanimoto scores is used for training. To achieve this, the spectrum pairs were picked by selecting the top 100 highest scoring library spectra for MS2Deepscore for a set of training spectra. This method for selecting spectrum pairs of spectra for training is similar to the workflow for running MS2Query and results in a relatively equal distribution of Tanimoto scores between these spectra, preventing a bias towards predicting lower Tanimoto scores.

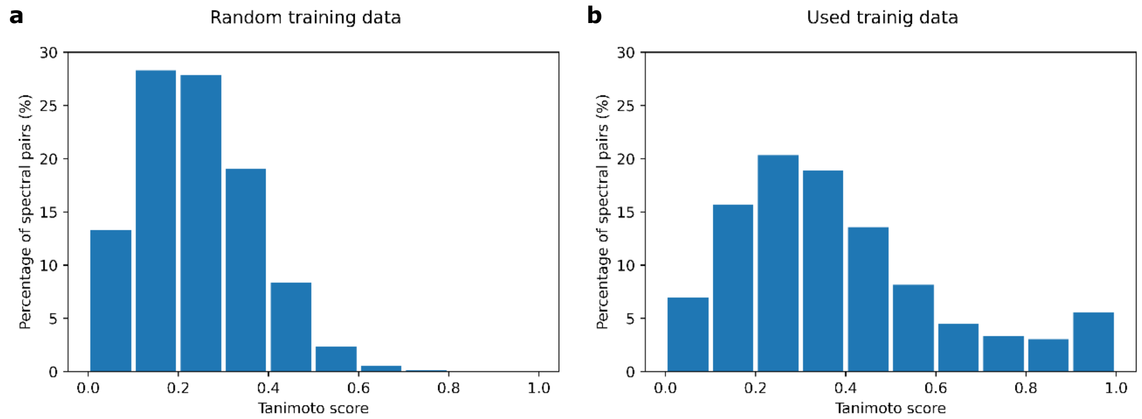


*Figure S5: **a**: Distribution of Tanimoto scores when selecting random spectral pairs. Spectral pairs are selected by making a pair between the training spectra and all InChiKeys in the library. **b**: Distribution of Tanimoto scores between the pairs of spectra used for training the random forest model. These spectral pairs were selected by calculating MS2Deepscore between each training spectrum and the reference library and making pairs with the 100 highest scoring library spectra.*

### S6. Additional analysis case study GNPS

The GNPS analogue search was run for case studies 1-3 using a mass window of 200 Da, a cosine score threshold of 0.6, minimum matched peaks set to 3, the ion mass tolerance set to 0.25 Da and a fragment Ion Mass Tolerance of 0.01 Da.

A molecular network was created with the Feature-Based Molecular Networking (FBMN) workflow (Nothias et al., 2020) on GNPS (https://gnps.ucsd.edu, (Wang et al., 2016). The feature quantification table and MS2 spectral summary file were uploaded to GNPS. The data were filtered by removing all $MS^2$ fragment ions within +/- 17 Da of the precursor m/z. $MS^2$ spectra were window filtered by choosing only the top 6 fragment ions in the +/- 50 Da window throughout the spectrum. The precursor ion mass tolerance was set to 0.02 Da and the $MS^2$ fragment ion tolerance to 0.02 Da. A molecular network was then created where edges were filtered to have a cosine score above 0.7 and more than 3 matched peaks. Further, edges between two nodes were kept in the network if and only if each of the nodes appeared in each other’s respective top 10 most similar nodes. Finally, the maximum size of a molecular family was set to 100, and the lowest scoring edges were removed from molecular families until the molecular family size was below this threshold.

The jobs can be publicly accessed at:

Blood plasma LTR analogue search:
https://gnps.ucsd.edu/ProteoSAFe/status.jsp?task=5d4577850dae44758da85c6ee3b77e89

Urine LTR analogue search:
https://gnps.ucsd.edu/ProteoSAFe/status.jsp?task=303ee7013d994074b27de8b86fd3bbad

Blood plasma NIST 1950 analogue search:
https://gnps.ucsd.edu/ProteoSAFe/status.jsp?task=6f0c60c689bb4facb64e96fac968ff0d

Anammox bacteria molecular networking:
https://gnps.ucsd.edu/ProteoSAFe/status.jsp?task=250044393bf44654ad72b7cebebba478

The results of the analogue searches are merged with the annotated csv files. Despite our efforts, it is challenging to judge for a case study what method performs better, due to the relatively low number of (tentatively) validated spectra and the fact that judging the quality of analogues is somewhat subjective. However, for case study 1 a relatively high number of case study spectra could be validated. In total, for 67 spectra, the precursor m/z and retention time could be linked to an in-house reference. Both the results predicted by GNPS and MS2Query were compared to the reference standards. The biggest difference was in the spectra that we were not able to validate, which makes it challenging to directly compare the performance of GNPS analogue search with GNPS for this case study.

*Table S4: Results of manual validation of results of MS2Query and GNPS analogue search for the NIST blood plasma case study.*

| | MS2Query | GNPS |
|---|---|---|
| Correct | 9 | 6 |
| Good analogue | 31 | 40 |
| Analogue | 16 | 11 |
| Bad analogue | 4 | 4 |
| Wrong | 1 | 0 |
| Unknown | 14 | 29 |
| Unannotated | 28 | 13 |
| Total | 103 | 103 |

### S7. Performance Negative Mode

The negative mode spectra were processed in the same way as described for the positive mode spectra in the materials and methods. For the negative mode less spectra are available that also cover less molecules in chemical space. The performance on an analogue search by MS2Query still has a higher accuracy compared to the modified cosine score and MS2Deepscore. However for searching for exact matches MS2Query performs less well than alternative methods. The performance of MS2Query is probably less well due to the fact that there is less training data available. With the increase of publicly available mass spectrometry data it is expected that the performance by MS2Query for negative mode spectra will also increase.

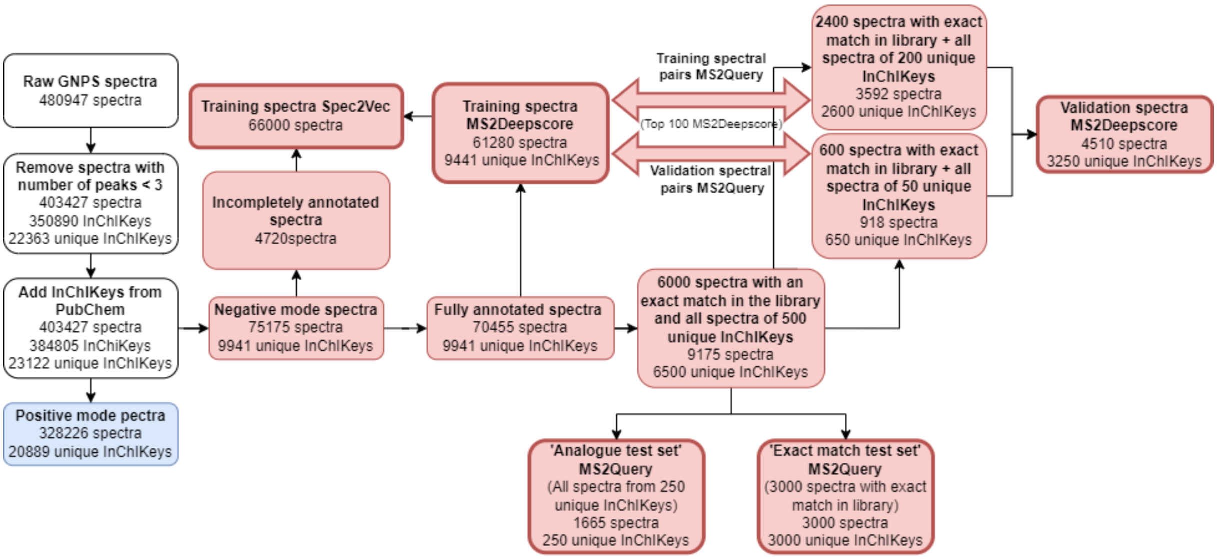


*Figure S6: Workflow for creating datasets used for training, validation and testing of MS2Deepscore, Spec2Vec and MS2Query for spectra in negative ionization mode.*

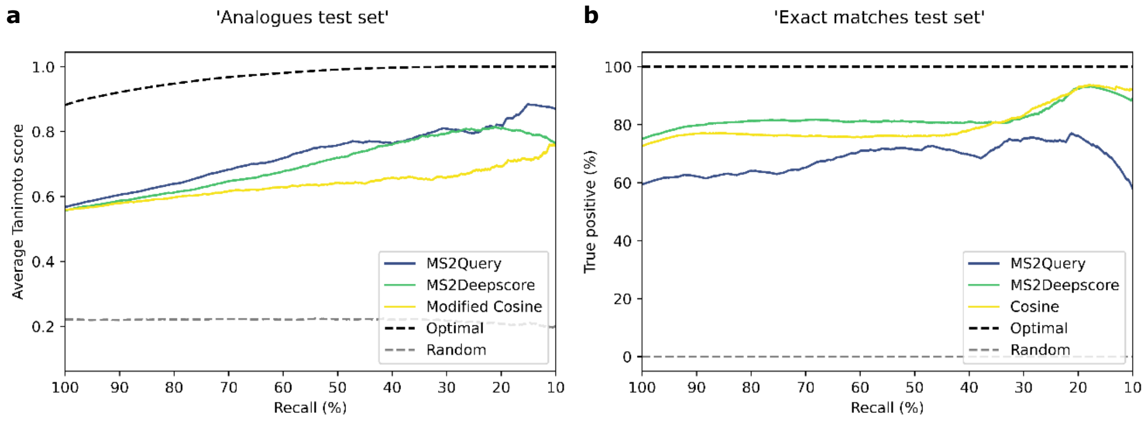


*Figure S7: Comparison of performance for MS2Query, MS2Deepscore and (modified) cosine score for a library and test sets of negative mode spectra. The threshold for MS2Query, MS2Deepscore and (modified) cosine is varied, resulting in different recalls.* ***a****: The 'analogues test set' is used with spectra that have no exact match in the library, therefore the best possible match is always an analogue. For MS2Deepscore and modified cosine score, library spectra are first filtered on a mass difference of 100 Da. The relationship between recall and accuracy is plotted. For each threshold the accuracy is measured by taking the average over the Tanimoto scores (chemical similarity) between the correct molecular structure and the predicted analogues.* ***b****: The 'exact match test set' of 3000 spectra is used, all these test spectra have at least 1 exact structural match in the reference library. For MS2Deepscore and modified cosine score, library spectra are first filtered on a mass difference of 0.25 Da, while MS2Query does not use any pre-filtering on mass difference, and uses the exact same settings as for the analogue search. The percentage of true positives is given, a match is marked as true positive if the first 14 characters of the InChiKeys are identical.*
